# Mitochondrial RNA processing in absence of tRNA punctuations in octocorals

**DOI:** 10.1101/103036

**Authors:** Gaurav G. Shimpi, Sergio Vargas, Angelo Poliseno, Wörheide Gert

## Abstract

**Background:** Mitogenome diversity is staggering among early branching animals with respect to size, gene density and content, gene orders, and number of tRNA genes, especially in cnidarians. This last point is of special interest as tRNA cleavage drives the maturation of mitochondrial mRNAs and is a primary mechanism for mt-RNA processing in animals. Mitochondrial RNA processing in non-bilaterian metazoans, some of which possess a single tRNA gene in their mitogenomes, is essentially unstudied despite its importance in understanding the evolution of mitochondrial transcription in animals.

**Results:** We characterized the mature mitochondrial mRNA transcripts in a species of the octocoral genus *Sinularia* (Alcyoniidae: Octocorallia), and defined precise boundaries of transcription units using different molecular methods. Most mt-mRNAs were polycistronic units containing two or three genes and 5’ and/or 3’ untranslated regions (UTRs) of varied length. The octocoral specific, mtDNA-encoded mismatch repair gene, *mtMutS*, was found to undergo alternative polyadenylation (APA), and exhibited differential expression of alternate transcripts suggesting a unique regulatory mechanism for this gene. In addition, a long noncoding RNA complementary to the ATP6 gene (lncATP6) potentially involved in antisense regulation was detected.

**Conclusions:** Mt-mRNA processing in octocorals bearing a single mt-tRNA is complex. Considering the variety of mitogenome arrangements known in cnidarians, and in general among non-bilaterian metazoans, our findings provide a first glimpse into the complex mtDNA transcription, mt-mRNA processing, and regulation among early branching animals and represents a first step towards understanding its functional and evolutionary implications.

## 1 Introduction

Two major evolutionary events occurred early in the animal history forging the majority of animals, as we know them today: the origin of their multicellularity, and the origin of bilateral symmetry. Multiple genomic changes accompanied these morphological transitions, and different genome sequencing projects give us a glimpse into these changes [1, 2]. Undoubtedly, these transitions also correlate with multiple changes in mitochondrial genome (mitogenome) architecture and organization [3]. The metazoan mitochondrial genome underwent reductive evolution, transferring most of its genome content to the nucleus [4, 5]. The majority of these alterations in mitogenome content include the loss of ribosomal proteins, and some tRNA genes, changes in the genetic code, disappearance of introns, and further compaction of mitochondrial DNA (mtDNA). As an aftermath, a quintessential animal mitochondrial genome harbors only 13 genes encoding essential energy pathway proteins, 2 ribosomal RNA genes and 22 transfer RNA genes. This composition is nearly invariable among bilaterians in terms of gene content [6]. However, alterations in mitogenome content, size and organization are more prominent and peculiar among non-bilaterian animals. The mitogenomes of non-bilaterian metazoan phyla comprise several novelties compared to canonical animal mitogenomes [7]. These include, the presence of group I and group II introns in sponges and scleractinians [8–12], additional protein coding genes and/or unknown ORFs and gene duplications in anthozoans [13–16], linear mitogenomes in calcisponges and medusozoans [17, 18], among other. Therefore, as a hotspot of mitochondrial genome diversity, early branching animals present a unique opportunity to understand the evolution of mitochondrial genome architectures as well as the fundamental processes governing its functionality and maintenance.

The significantly reduced but extremely crucial repertoire of genes present in animal mitogenomes is fundamental to its molecular and cellular functions, and to gain a deeper understanding of these processes it is essential to understand the expression and processing of mitochondrial gene transcripts. The majority of information available so far on the mitogenome transcription originates from bilaterian members of the animal kingdom [19–22]. In this regard, the canonical vertebrate mitogenome is known to transcribe symmetrically as polycistronic precursors spanning the entire heavy (H-) and light (L-) strands [23]. The 22 tRNAs interspersed throughout the mitogenome serve as punctuation marks that are recognized and cleaved at 5’ and 3’ ends by the mitochondrial RNase P and RNase Z, respectively [24, 25]. The genes within these precursors are simultaneously liberated for maturation following this mt-tRNA processing step. Consequently, in most bilaterian metazoans, most mature mitochondrial mRNAs are monocistronic, with *ATP8-ATP6* and *ND4L-ND4*, which are known to exist as bicistronic elements, as the only exceptions. Finally, all messengers end with the post-transcriptional addition of 40-45 adenosines for maturation, which also completes the stop codon at 3’ end of mRNA in most cases [26]. Bilaterian mt-mRNAs are either essentially devoid of the untranslated regions (UTRs) or tend to have very short UTRs consisting of 1-2 nucleotides flanking the mature mRNAs; a few notable exceptions such as *COI*, *COII*, *ND5* and *ND6* genes, which possess slightly longer 3’ UTRs complementary to the genes on opposite strand [23–27]. This transcription model is, however, based on the study of a small number of bilaterians [19–22]. Recent studies on medusozoan members possessing linear mitogenomes do provide some insights into the mt-transcription in non-bilaterians [28, 29]. However, a detailed exploration of mitochondrial RNA processing and characterization of UTRs is still lacking for most non-bilaterian animals, including anthozoans with circular mitogenomes more similar to the stereotypical animal mitochondrial genomes.

Among non-bilaterian metazoans, octocorals (Octocorallia: Anthozoa) are unique due to their atypical mitochondrial genomes. As many as five different gene arrangements have been reported among the octocorals studied so far [30–33], all with an exceptionally reduced complement of transfer RNAs (i.e. a single tRNAMet gene) and the presence of an additional gene, a mismatch repair gene (*mtMutS*) [13, 34], closely related to the non-eukaryotic *MutS7* lineage from epsilon-proteobacteria or DNA viruses [35]. This gene has been predicted to have a self-contained mismatch DNA repair function [35], and it has been speculated to play a role in the slow rate of mtDNA evolution observed among octocorals [36, 37], and to be responsible for various genome rearrangements through intramolecular recombination [31].Considering the evolutionary trend towards a reduced mitogenome in Metazoa [38], the occurrence of such a large gene, such as the *mtMutS*, occupying nearly 16% of the octocoral mitogenome is somewhat surprising. The presence of a *mtMutS* mRNA transcript suggested its availability for translation [35]. However, 20 years after its discovery [13] and despite of being extensively used for phylogenetic studies of octocorals [39], a thorough understanding of its transcriptional processing and maturation, and in general of its biology is lacking.

Octocoral mitogenomes exhibit five different gene arrangements, all containing a single gene for tRNA. Cleavage of this tRNA from the precursor polycistronic RNA would only result in linearization of precursors. The way in which the individual mitochondrial gene mRNAs are released for maturation from the long polycistronic precursor remains to be determined. In absence of knowledge on the precise boundaries of the mitochondrial mRNA in octocorals, despite all the novelties they confer, the understanding of the biology and evolution of animal mitochondria remain incomplete. Here we characterize the mitogenome transcription of an early branching non-bilaterian metazoan, the octocoral *Sinularia* cf. *cruciata*. (Alcyoniidae: Octocorallia). We describe the 5’ and 3’ boundaries and UTRs of mature mitochondrial mRNAs and characterize the transcription of the *mtMutS* gene. Our results provide the first glimpse to the unique features and complexity of the mitochondrial transcriptome in non-bilaterians.

## 2 Materials and Methods

### Abbreviations

*COI*: Cytochrome oxidase, subunit I; *COII*: Cytochrome oxidase, subunit II; *ATP6*: ATP synthase, subunit 6; *ATP8*: ATP synthase, subunit 8; *ND5*: NADH dehydrogenase, subunit 5; *ND4*: NADH dehydrogenase, subunit 4; *ND4L*: NADH dehydrogenase, subunit 4L; *ND6*: NADH dehydrogenase, subunit 6; *CytB*: Cytochrome b; *ND1*: NADH dehydrogenase, subunit 1; *ND2*: NADH dehydrogenase, subunit 2; *ND3*: NADH dehydrogenase, subunit 3; *COIII*: cytochrome oxidase, subunit III; 12S: Small subunit of mitochondrial ribosomal RNA gene; 16S: Large subunit of mitochondrial ribosomal RNA gene.

### Specimens

Coral colonies were obtained from a commercial source and maintained in a closed circuit seawater aquarium at Molecular Geo- and Palaeobiology lab, LMU, Munich. Two species of the genus *Sinularia* were used in this study. *Sinularia* cf. *cruciata* (Museum Voucher Code: GW1725) was utilized for the majority of the experiments, whereas additionally, a second *Sinularia* sp. (Museum Voucher Code: GW2911) was used for RT-PCR screening, RHAPA and antisense mRNA detection (see below). All the references to the nucleotide positions refer to the full mitochondrial genome of *Sinularia* cf. *cruciata* (GenBank accession KY462727), which was sequenced completely (details are provided in Additional file 1).

### Total RNA extraction and cDNA synthesis

TRIzol reagent (Invitrogen, USA) was utilized for the extraction of total RNA as per the manufacture’s instructions. RNA was dissolved in 100 µl DEPC treated water and contaminating DNA was eliminated from RNA extracts by performing a DNase (RQ1 RNase-free DNase, Promega, USA) treatment at 37 °C for 30 min. Treated RNA was purified after inactivation of the DNase and its purity was determined using a Nanodrop ND-1000 spectrophotometer (Thermo Fisher Scientific, USA). RNA samples with absorbance at OD260/280 and OD260/230 ratios ∼2.0 were used for further analysis. RNA integrity was also verified by 1% agarose gel electrophoresis as well as using a Bioanalyzer (Agilent Inc.). RNA extracts with a RIN value ≥7.5 were used for cDNA synthesis (data not shown); these extracts were stored at −80°C until use.

### RNA-Seq and read mapping

RNA-Seq reads for *Gorgonia ventalina* (SRR935078-SRR935087) and *Corallium rubrum* (SRR1552943-SRR1552945 and SRR1553369) were downloaded from NCBI’s Short Read Archive, imported in Geneious and mapped against the mitochondrial genomes of *Pseudopterogorgia bipinnata* (NC_008157), in the case of *G. ventalina*, or *Corallium rubrum* (AB700136). In the case of *Sinularia* cf. *cruciata* ca. 28*x*10^6^ 50bp pairs of reads were sequenced, imported in Geneious 8.1.8 (Biomatters) [40] and mapped to the mitochondrial genome we sequenced for this species. The mapping was done using a Low Sensitivity strategy that avoids remapping reads to previously build contigs based on the previous mapping rounds. The mapping results were screened in Geneious to find gaps in coverage flanking putative transcription units. Additionally, we screened the mapped reads to assess whether read pairs spanning adjacent genes could be found among the sequences. These reads, i.e. read-pairs linking adjacent genes and spanning intergenic regions, were taken as evidence of collinearity.

### Reverse-transcription PCR (RT-PCR)

RNA extracts were PCR controlled in order to detect amplifiable levels of small DNA fragments. Only RNA extracts devoid of any amplification were used in RT-PCR experiments. For each sample, ∼1 µg of total RNA was reverse transcribed in 20 µl reactions using the ProtoScript II First Strand cDNA Synthesis Kit (New England Biolabs, USA) with an anchored oligo-(dT) primer and following the manufacture’s instructions. RT-PCR sequencing primers were designed using the sequenced mitochondrial genomic sequence of *Sinularia* cf. *cruciata*, GenBank accession KY462727. Screening for the presence of the polycistronic mRNAs was done using the primers enlisted in Table 1.

**Table 1:** RT-PCR screening for mature mRNA transcripts spanning two or more genes. Primer codes: first part indicates gene and number indicates 5’ end of primer corresponding to *S*. cf. *cruciata* mitogenome; + represents positive amplification, − represents negative amplification. Asterisk (*) indicates bps into the gene downstream to the igr

| Sr. No. | Primer Pair (Forward/Reverse) | Regions | Length (bp) | igr + Gene* | RT-PCR DNA | cDNA |
| --- | --- | --- | --- | --- | --- | --- |
| 1 | C2-18519/C1-839 | *COII-COI* | 1051 | 951 | + | – |
| 2 | C1-743/12S-2240 | *COI-12S* | 1498 | 658 | + | – |
| 3 | N1-3534/CB-4128 | *ND1-CytB* | 595 | 479 | + | – |
| 4 | CB-4539/N6-4995 | *CytB-ND6* | 447 | 142 | + | + |
| 5 | CB-4539/N3-5625 | *CytB-ND6-ND3* | 1064 | *ND6*+171 | + | – |
| 6 | N6-4974/N3-5625 | *ND6-ND3* | 641 | 171 | + | + |
| 7 | N6-4974/N3-5820 | *ND6-ND3* | 859 | 389 | + | – |
| 8 | N6-4974/N4L-6120 | *ND6-ND3-ND4L* | 1159 | *ND3*+291 | + | – |
| 9 | N3-5499/M-6780 | *ND3-ND4L-mtMutS* | 1282 | *ND4L*+638 | + | + |
| 10 | N3-5499/M-7199 | *ND3-ND4L-mtMutS* | 1701 | *ND4L*+1057 | + | + |
| 11 | N3-5499/M-8286 | *ND3-ND4L-mtMutS* | 2788 | *ND4L*+2456 | + | + |
| 12 | N3-5499/M-8826 | *ND3-ND4L-mtMutS* | 3328 | *ND4L*+2684 | + | + |
| 13 | M-8714/16S-9396 | *mtMutS-16S* | 683 | 259 | + | + |
| 14 | 16S-10954/N2-11753 | *16S-ND2* | 800 | 639 | + | – |
| 15 | N2-12177/N5-12612 | *ND2-ND5* | 448 | 106 | + | + |
| 16 | N5-12961/N4-15314 | *ND5-ND4* | 2354 | 990 | + | + |
| 17 | N4-14807/C3-16468 | *ND4-COIII* | 1662 | 598 | + | – |
| 18 | N4-15714/A6-17048 | *ND4-COIII-ATP6* | 1335 | *COIII*+225 | + | – |

### Analysis of 5’ and 3’ ends

Two different approaches were used to determine the transcript ends of mature mitochondrial mRNA species.

#### Circularized RT-PCR (cRT-PCR)

Isolation of mRNA from total RNA was performed using Dynabeads mRNA magnetic beads (Invitrogen). 100 ng of polyA-selected mRNA as well as total RNA were circularized using T4 RNA ligase I (New England Biolabs) following the manufacture’s protocol. The circularized RNA was purified and used for cRT-PCR and 5’/3’ end screening using the method described in [41]. Briefly, cDNA synthesis was performed as described above using gene-specific reverse primers binding near the 5’ end of linear RNA resulting in production of first strand that contains 5’ end, the ligation site, and the 3’ end of the molecule. These first strands were subjected to PCR amplification using specific primer pairs (see Table 2 for primer details).

#### Rapid amplification of cDNA ends (5’and 3’ RACE)

First strand synthesis was performed to obtain the template for 5’ and 3’ RACE PCRs using SMART RACE cDNA Amplification Kit (CLONETECH Inc.) following the supplier’s protocol. Approximately ~1 Âţg of total RNA was used to obtain two separate pools of 5’-RACE-Ready cDNA and 3’-RACE-Ready cDNA. RACE PCR reactions were performed using different gene-specific primers paired with adaptors primers as per the supplier’s instructions (see Table 2).

**Table 2:**
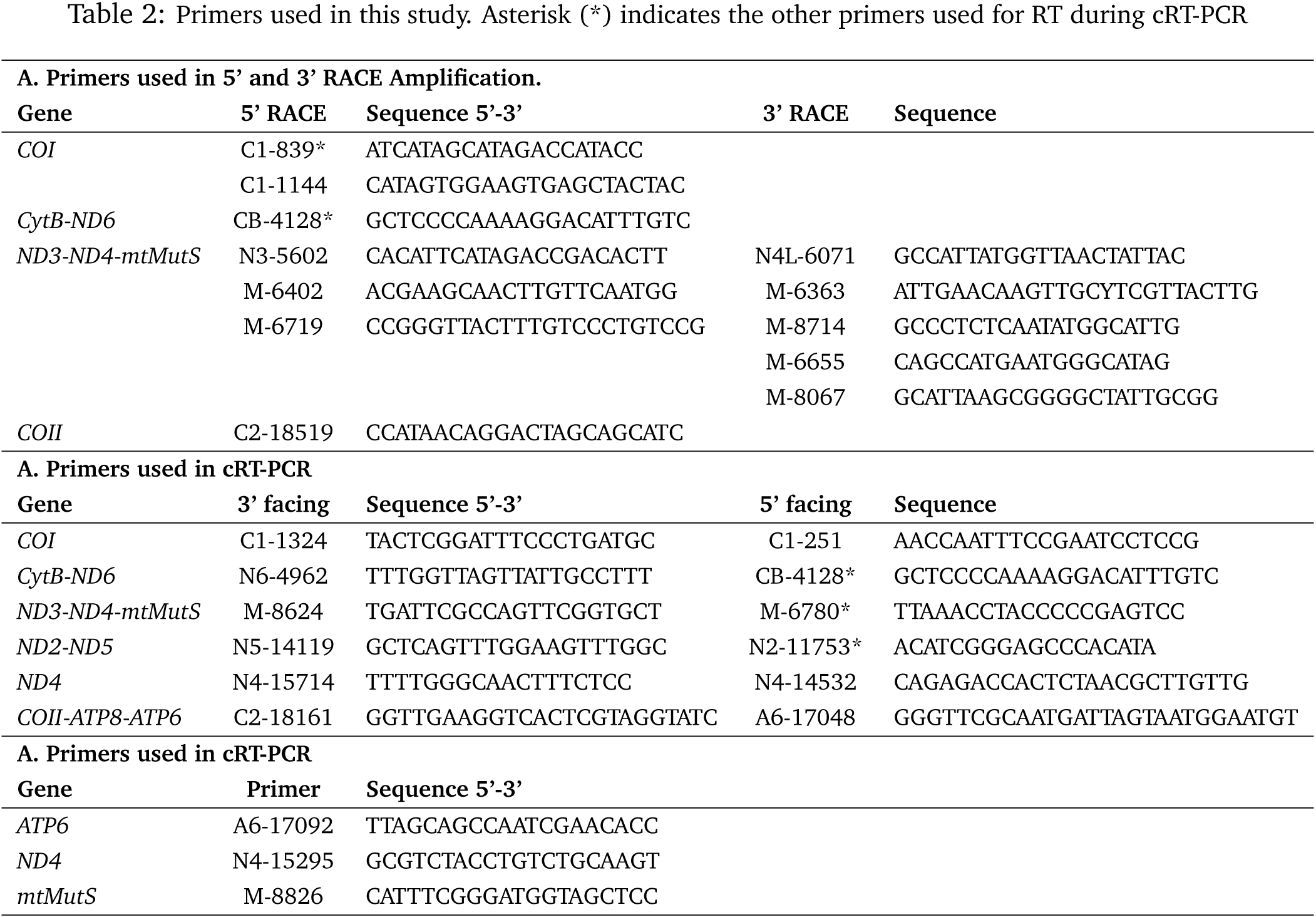
Primers used in this study. Asterisk (*) indicates the other primers used for RT during cRT-PCR

### Cloning, sequencing and sequence analysis of amplified products

Amplified products were either extracted from 1% agarose gels or purified using the NucleoSpin Gel and PCR Purification Kit (MACHEREY-NAGEL, Germany) and cloned using a TOPO TA Cloning Kit (Invitrogen, USA). The clones obtained were PCR amplified, precipitated and sequenced using ABI BigDye v3.1 sequencing chemistry on an ABI 3730 DNA Analyzer Sequencing instrument. Sequences obtained were analyzed and aligned to the mitogenomes of S. *piculiaris* (NC_018379) and the sequenced S. cf. cruciata (Accession No. KY462727) using Geneious 6.1.6 software (Biomatters) [40].

### Detection and quantification of alternative polyadenylation (APA)

RNase H alternative polyadenylation assay or RHAPA [42] was employed to determine and quantify alternative polyadenylation of the *mtMutS* mRNA transcripts. For the first time, we coupled this assay with quantitative real-time PCR technique (qPCR), which allows for the accurate estimation of the abundance of the alternative transcripts. The oligonucleotide 5’-CATTTCGGGATGGTAGCTCC-3’ was used to remove the 3’ end of the polyadenylated complete *mtMutS* messenger. This primer hybridizes to the *mtMutS* mRNA between positions 8807-8826. After hybridization, the DNA-RNA hybrids were digested with RNase H and the resulting mRNA was purified using RNA Clean & Concentrator kit (Zymo Research) and reverse transcribed using oligo(dT) as described above. Only alternatively polyadenylated *mtMutS* forms should be present in the resulting cDNA after RNase H digestion. A control RT-PCR using primers binding to the adjacent regions of the RNase H digested site ensured the successful digestion of the 3’ including and poly(A) tail of the mature, full *mtMutS* mRNA species. Afterwards, a quantitative real-time PCR (qPCR) assay was performed to determine the abundance-levels of the transcripts using primers binding upstream and downstream of digested mRNA region. For primer details see Additional file 4, Table 2.

### Strand-Specific RT-PCR

Strand-specific RT-PCR was performed as described previously [43] for the detection of antisense RNA transcripts of the *ATP6* gene in both *Sinularia* species. In earlier, RACE experiment, we observed *ATP6* clones containing a RACE adaptor ligated at 3’ end of 5’ RACE-ready cDNA. This suggested the presence of an antisense transcript for this gene, as it has also been noted previously in porcine brain [44]. Two antisense strand-specific primers were used for cDNA synthesis (AAR1. 5’-TTACTCCTACTGCCCATATTG-3’ and AAR2. 5’-TGTAGTTCGGATAATTGGGGG-3’), whereas a sense strand-specific primer (SAF. 5’-TTAGCAGCCAATCGAACACC-3’) and an anchored oligo(dT) were employed separately for first strand synthesis. For RT-PCR, the primer AAR1-SAF pair was used.

### Quantitative Real-time RT-PCR (qPCR) and data analysis

The Rotor-Gene Q 2plex system (QIAGEN) was utilized for qPCR experiment. The KAPA SYBR FAST universal mastermix (Peqlab) was used in 15μl reactions containing 1 μl diluted cDNA, 7.5 μl 2X mastermix, and 250 to 400 nM of each primer. A two-step qPCR including an initial denaturation step of 3 min at 95 °C followed by 40 cycles of 95 °C for 10 s and 60 °C for 20 s was performed. A nontemplate control was always included in each assay and melting curve analysis was performed at the end of each qPCR to confirm amplification specificity. In addition, amplification products were also checked by agarose gel electrophoresis after each assay.

Fluorescence data obtained after qPCR was analyzed using LinRegPCR, which determines Cq values and PCR efficiencies [45]. These values were further used to analyze mitochondrial expression; statistical tests were performed using REST 2009 (QIAGEN) as described previously [46].

## 3 Results

### The mitogenome of *Sinularia* cf. *cruciata*

The complete mitogenome of *Sinularia* cf. *cruciata* was 18,730 bp in length and included, similar to other octocorals, 14 protein-coding genes (PCGs) (*ATP6*, *ATP8*, *COI*-III, *CytB*, *ND1*-6, *ND4L* and *mtMutS*), two ribosomal RNAs (12S and 16S) and a single transfer RNA (tRNAMet). Most PCGs and the two rRNA genes were encoded on the H-strand. *ATP6*, *ATP8*, *COII*, *COIII* and tRNAMet were encoded on the L-strand. Gene order was consistent with that of other octocorals with mitogenome arrangement “A” [31]. Base composition of the mitogenome was A, 30.2%; C, 16.5%; G, 19.3%; T, 33.9% and G+C, 35.8%. Among the 14 PCGs, the *mtMutS* (2982 bp) was the longest and *ATP8* (216 bp) the shortest. All PCGs had ATG as the start codon, while the stop codons TAA and TAG were predominant among PCGs; *COI* was an exception, having an incomplete termination codon (T) (see Additional file 1, Table 1). Except for *ND2* and *ND4* (13 bp overlap), the remaining genes were separated by intergenic regions (IGRs) of different lengths (see Additional file 1, Table 1 and Figure 1A). The longest IGR (112 bp) was found between *COII-COI*, while the shortest (4 bp) was located between 12S-*ND1*. The mitogenome of *Sinularia* cf. *cruciata* was 12 bp shorter than that of *Sinularia peculiaris* and the two species had same base composition and GC content. Sequence variability within the two *Sinularia* species was 2.31%, excluding nucleotide ambiguities (0.05%) and gaps (0.16%). The most variable genes were *mtMutS*, *ND2*, *ND5*, *ND4* and *COII* (Additional file 1, Figure 1B).

**Figure 1:**
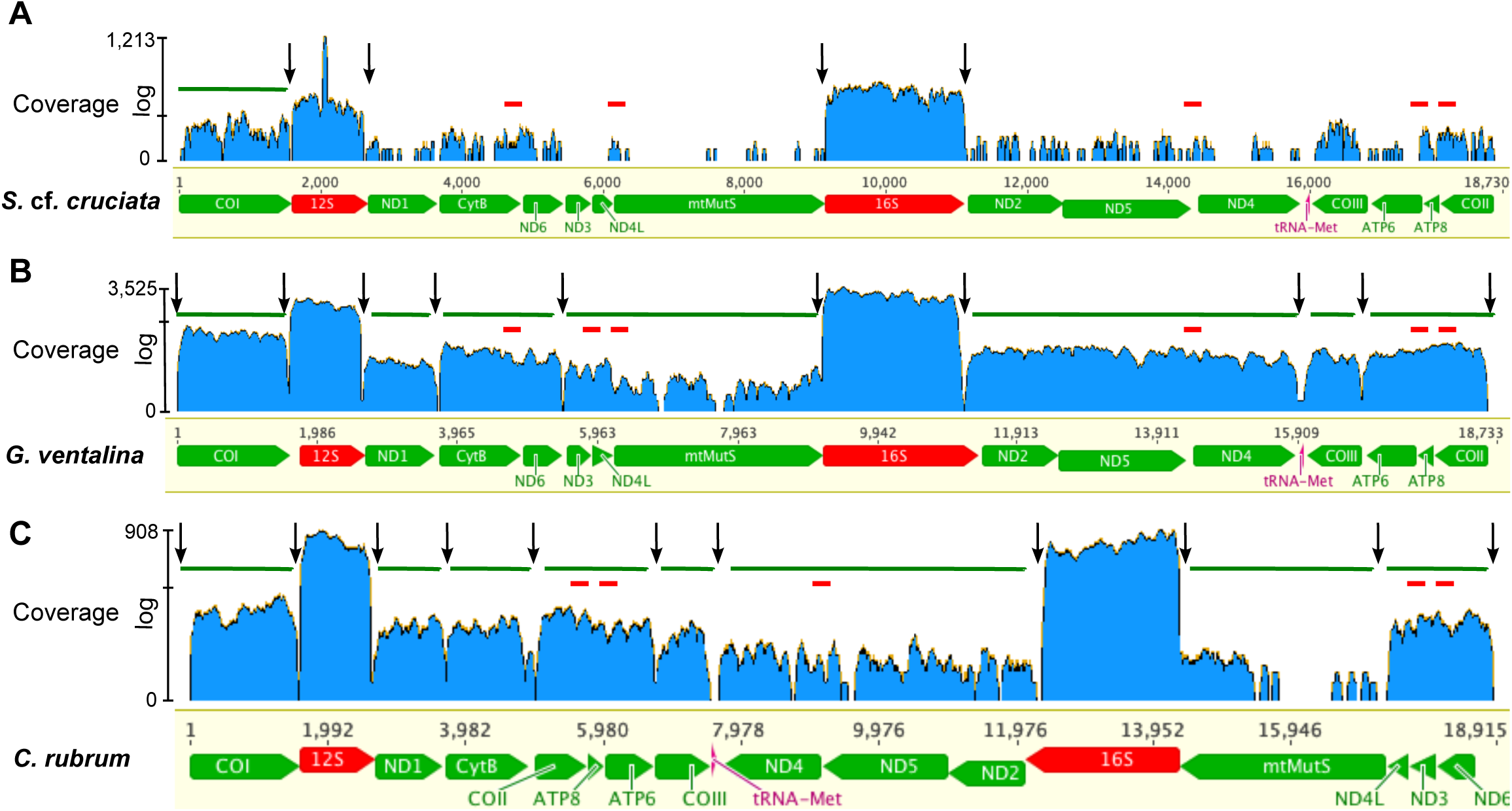
RNA-Seq (log scale) coverage of the mitochondrial genome of octocorals (A) *Sinularia* cf. *cruciata*, (B) *Gorgonia ventalina*, and (C) *Corallium rubrum*. The X-axis represents positions of the mitochondrial genes. Green and red blocks represent PCGs and rRNAs, respectively. Y-axis shows the log of number of reads mapped at each nucleotide. Black arrows indicate abrupt drop in coverage. Green bars above coverage represent putative transcriptional units of PCGs, whereas small red bars below show presence of reads for IGRs.

### Recovering mitogenomic transcripts from RNA-Seq data of S. cf. *cruciata* and other octocorals

In order to understand mtDNA expression and processing in octocorals, an RNA-Seq library of *Sinularia* cf. *cruciata* was screened for mitochondria-mapping reads (hereafter mt-reads). A total of 4,153 reads out of 26M pairs were mapped to the sequenced *S.* cf. *cruciata* mitogenome. This resulted in a partial mitogenome assembly covering 62.8% of its length and leaving 37.2% of the genome uncovered. With the exception of 12S and 16S rRNAs, which exhibited very high coverage, none of the PCGs present in the mitogenome were completely covered (Figure 1). Despite low coverage, reads spanning IGRs were detected pointing towards the collinear expression of *CytB-ND6*, *ND4L-mtMutS*, *ND5-ND4*, *ATP8-ATP6*, and *COII-ATP8*.

Additionally, RNA-Seq data from two published octocoral transcriptome studies were screened for mitochondrial reads. These included *Gorgonia ventalina* (Genome arrangement A; SRX177804-5) and *Corallium rubrum* (Genome arrangement C; SRX675792) [31, 47–49]. Hence, they provided us with the opportunity to explore mt-transcription in octocorals with different mitogenome arrangements. In case of *G. ventalina*, 55,783 out of 302M reads were mapped to the *Pseudopterogorgia bipinnata* mitogenome (NC_008157), a species closely related to *G. ventalina*. Almost the entire mitogenome was covered except for 2.1% uncovered sequence data, which mainly included IGRs between putative transcription units and a region of the *mtMutS* gene. Based on the observation of read-pairs spanning IGRs, the entire L-strand genes, namely *COII*, *ATP6*, *ATP8*, *COIII*, were detected as a collinear unit, whereas, for the H- strand the *COI*-12S-*ND1*, *CytB-ND6*, *ND3-ND4-mtMutS*, *ND2-ND5-ND4* were detected as collinear transcriptional units. However, as judged by sudden drop in coverage at IGRs between genes and the difference in expression levels of *COI* and 12S as well as 12S and *ND1* it is likely that these genes are actually monocistronic units and that the detected collinearity results from sequencing of low abundance premature RNA or unprocessed intermediates of these genes.

In the case of *C. rubrum*, 17,126 out of 241M reads were mapped to its mitogenome (AB700136). 8.8% (1,661 nt) data was missing. *COI* was present as a single transcriptional unit whereas other PCGs were observed to occur as a collinear units as follows: *ND1-CytB*, *COII-ATP8-ATP6-COIII*, *ND6-ND3-ND4L*, and *ND2-ND5-ND4* (with low coverage). We were unable to assign *mtMutS* to any of the transcription units due to small number of reads mapped to it. (see Figure 2 for a proposed scheme of mitogenome expression for arrangement “A” and “C”).

**Figure 2:**
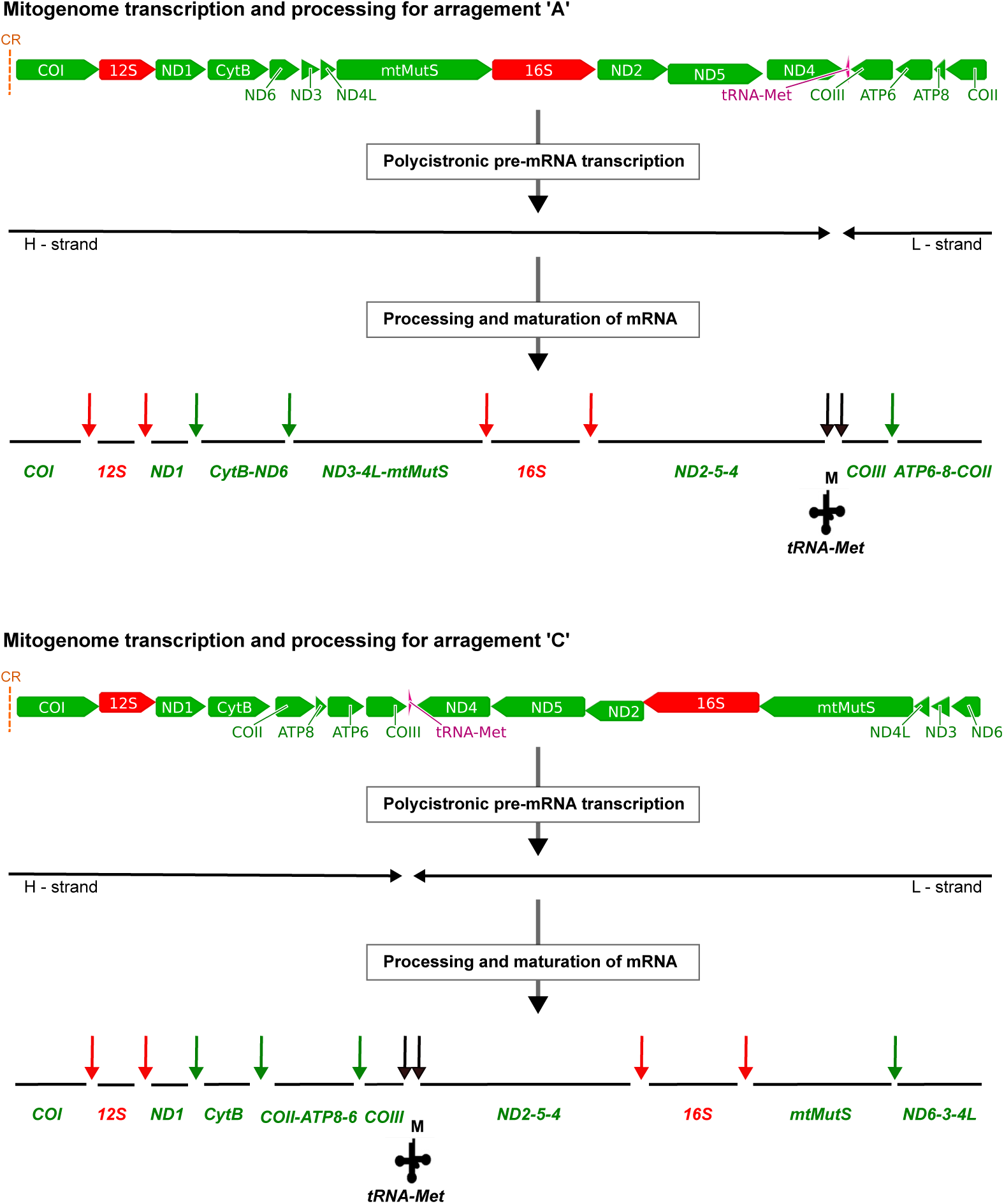
Predicted model of mt-mRNA processing for mitogenome arrangements ‘A’ and ‘C’ in octocorals. The vertical dotted orange lines are the control regions (CR). The black horizontal arrows are two polycistronic pre-mRNA transcripts each. Black vertical arrows indicate excision of tRNA as per “tRNA punctuation model”. Red and green vertical arrows represent additional excision sites and mature mitochondrial mRNA transcripts predicted based on our data.

Independently of the sequencing depth, in all three analyzed transcriptomes 12S and 16S were the most abundant transcripts (Additional file 2). Moreover, we observed that none of the three analyzed transcriptomes contained reads to cover the complete *mtMutS* gene transcripts. No RNA-Seq reads could be mapped to the IGR region between *COII-COI* and *ND6-COI* in *S.* cf. *cruciata* / *G. ventalina*, and *C. rubrum*, respectively. These regions fold into a stable stem-loop structure with a 33bp conserved motif in all octocorals studied here (Additional file 3), and represent an inversion of polarities, thus they could function as control regions (CR)/ origin of replication (OriH) in octocorals with “A” and “C” type mitogenome.

### RT-PCR corroborates the presence of mature polycistronic mRNA transcripts in the mitochondrial transcriptome of S. cf. *cruciata*

Depending on the sequencing depth of the RNA-Seq libraries, the RNA-Seq data may contain immature/unprocessed precursor RNA that could lead to the detection of false polycistronic mRNAs. In order to verify the presence of genuine mature polycistronic mRNA transcripts, we conducted RT-PCR experiment using primers binding substantially up/downstream from the start/stop codons of putative consecutive genes and amplifying their IGRs. Using DNA as a template, amplification was observed using all primer pairs screened (Table 1). Using cDNA synthesized with an anchored oligo(dT) primer as a PCR template, bicistronic and tricistronic transcripts corresponding to the *CytB-ND6*, *ND2-ND5*, and *ND3-ND4L-mtMutS*, *COII-ATP8-ATP6*, respectively, were detected corroborating the result obtained from the analysis of RNA-Seq data suggesting that these regions are polycistronic transcription units.

### UTR mapping of the mitochondrial protein coding genes in S. cf. *cruciata*

The UTRs of several mitochondrial mature transcriptional units were mapped using 5’/3’RACE and circularized RT-PCR. The *COI* mRNA has a 4bp 5’UTR upstream to this start codon. The 3’end of this gene, which lacks proper stop codon could not be deduced with enough certainty, but is likely that the partial stop codon is completed by polyadenylation yielding a monocistronic unit, in agreement with the transcriptome and RT-PCR results, which suggested a monocistronic nature of *COI*.

For *CytB* three different 5’ ends were detected. One of these 5’ ends initiated exactly two codons (6bp) downstream (position 3683) from the annotated start, without any 5’UTR. The other two were downstream from this 5’ end at positions 3926 and 3970. The first two messengers were detected using both RACE and cRT-PCR, whereas the third one was observed only with cRT-PCR. No 3’ end could be detected for *CytB* mRNA further supporting its co-expression with *ND6* in a bicistronic unit. Moreover, a polyadenylated *ND6* mRNA 3âᾸŹ end was detected with an end was detected with an 8bp 3’UTR.

Solely based on cRT-PCR, the mature mRNA ends were detected for the *ND2-ND5-ND4* tricistronic unit. The 5’ consists of a single base before the start codon at position 11173 (not 11146 as it is annotated in GenBank). A 44bp long 3’UTR was found after the stop codon at position 15868.

The analysis of transcriptomic data indicated the presence of a *COII-ATP8-ATP6* tricistronic mRNA. RT-PCR results and end mapping corroborated this observation. *COII* mRNA was found to be flanked by a 3bp 5’UTR, whereas an 83bp long 3’UTR was detected after the stop codon of *ATP6*. The precise ends of the protein coding genes *ND1* and *COIII*, which based on transcriptome and RT-PCR results, likely are monocistronic messages could not be determined. For more details on the UTRs of the mature mt-mRNAs of *S.* cf. *cruciata* see Figure 3.

**Figure 3:**
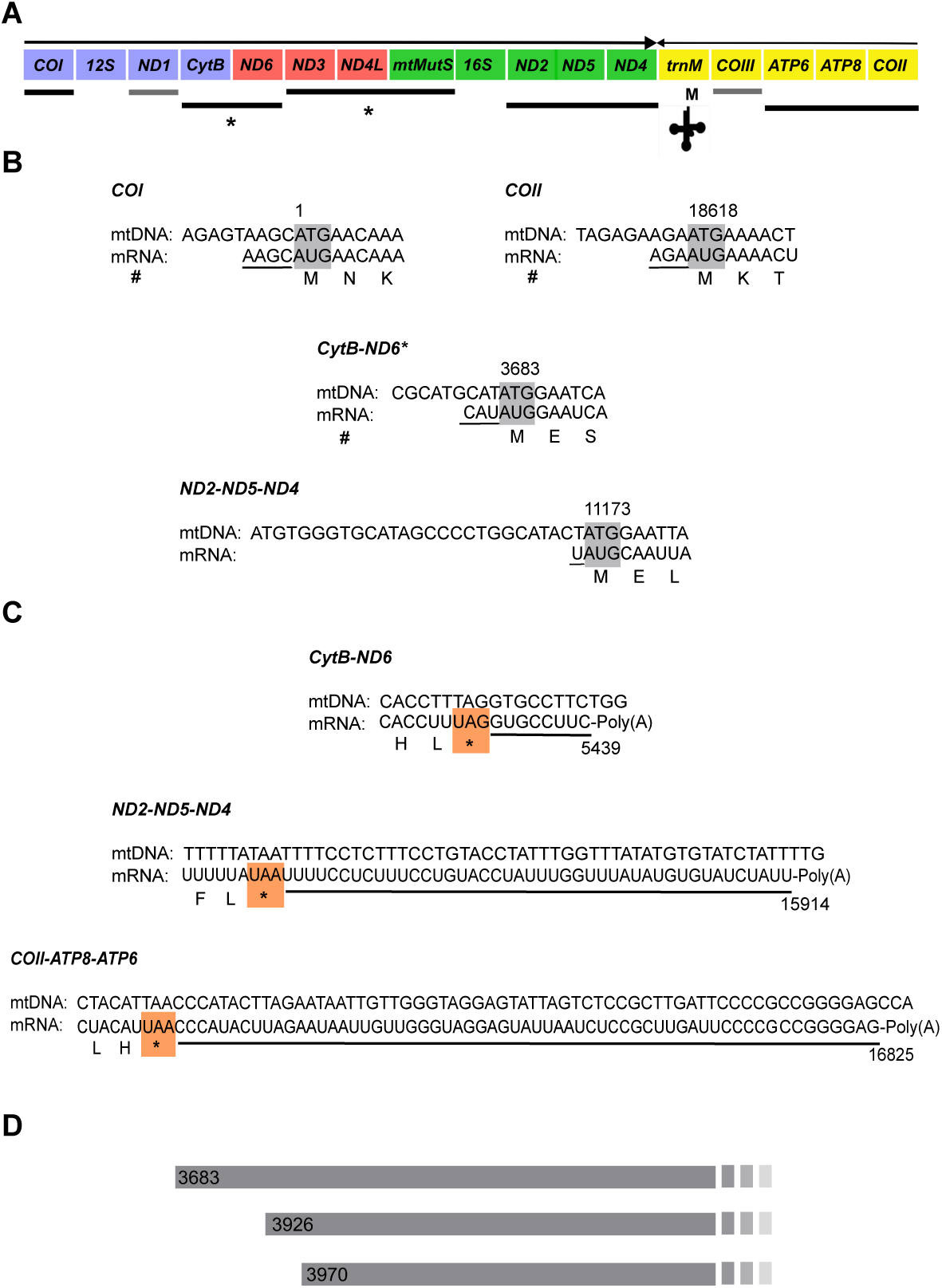
Mapped 5’ and 3’ ends of mature mitochondrial mRNAs. (A) Schematic representation of *Sinularia* mito-transcriptome arrangement. Arrows above show transcription orientation. Lines below denote the transcription units (mono- and polycistronic transcripts). Black lines below indicate transcription units for which one or both the ends are known, whereas grey lines (below *ND1* and *COIII*) indicate units for which ends remained unknown. Asterisk (*) shows the transcription units for which alternate ends were detected. (B) Summary of 5’ end mapping for mt-mtRNAs. The 5’ UTR regions are underlined. Shaded boxes depict start codons. # indicates detection using both, RACE and cRT-PCR methods. Nucleotide positions of the first base of the start codons are indicated. (C) Summary of 3’ end mapping for mt-mtRNAs. The 3’ UTR regions are underlined. Colored boxes highlight stop codons. Nucleotide positions of the last base of stop codons are indicated. (D) Alternative starting positions (5’ ends) of *CytB-ND6* mRNA.

### The *mtMutS* is alternatively transcribed and its transcripts are differentially expressed

For the *mtMutS*, 5’-RACE mapped a 31bp 5’ UTR upstream of the *ND3* start codon further corroborating that this gene is transcribed as a tricistronic unit together with *ND3* and *ND4L* (Figure 4A and B). Using several *mtMutS*-specific primers we were unable to detect any alternate 5’ end using RACE PCR. However, using circularized RT-PCR an internal *mtMutS* 5’ end was detected 387bp downstream (position 6542) of the annotated *mtMutS* start codon. The 3’ end for this particular transcript was at a position 8950, which is 188bp upstream to the annotated stop codon. This transcript ended with UAA and is clearly followed by a polyA-tail. However, this stop codon is not in frame.

Somewhat surprisingly, at least six different 3’ ends were detected for the *mtMutS* gene mRNA using RACE and cRT-PCR. These were found to end at position 6746, 6771, 6911, 8761, 8950 (as described above), and 8977 besides the annotated in-frame stop codon at position-9135 (Figure 4C). These messages are polyadenylated and end with GAA or UAA. Notably, none of these end-codons are in-frame. RT-PCR screening confirmed the presence of all possible transcript variants described above (see Additional file 4 Table 1 for primer details).

**Figure 4:**
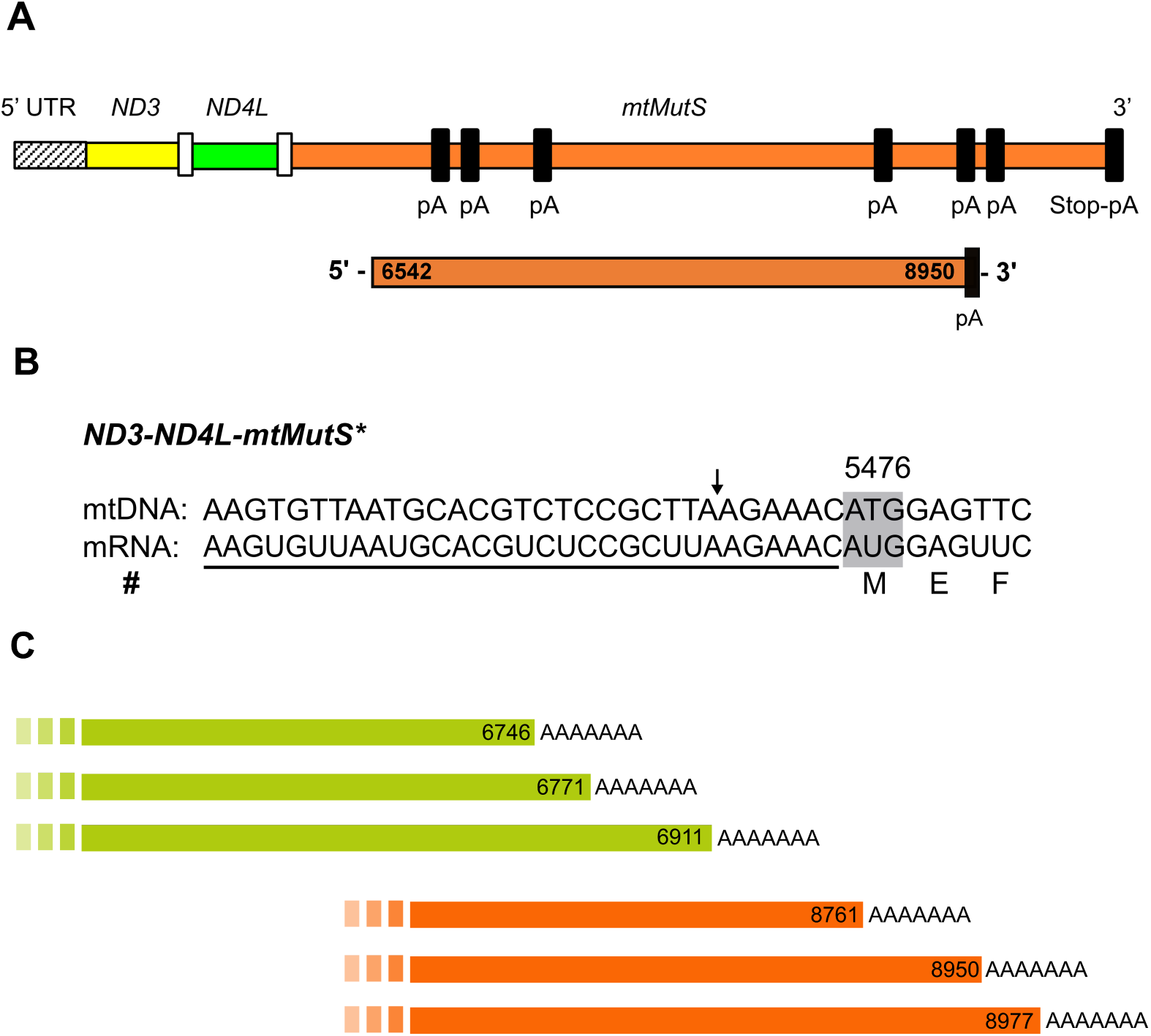
The *mtMutS* mRNA transcripts. (A) Schematic of the *mtMutS* gene as a tricistronic transcription unit with different poly(A) tail positions (not to scale) shown as dark blocks. pA=poly(A)-tail. Below is the internal *mtMutS* transcript. (B) The 5’ end of *ND3-ND4L-mtMutS* tricistronic transcript. The 5’ UTR region is underlined. Shaded box indicate the start codon. The arrow above indicates 11bp deletion in *S.* cf. *cruciata* compared to *S. piculiaris*. # indicates detection using both, RACE and cRT-PCR methods. (C) Alternatively polyadenylated *mtMutS* mRNAs; the position of the poly(A) start is indicated.

Further confirmation of alternative polyadenylation of *mtMutS* messengers in the normal mt-mRNA pool was attained using RHAPA (RNase H alternative polyadenylation assay). The *mtMutS* transcripts were differentially expressed. The central region, which includes a partial domain III of *mtMutS*, was 6.35±0.3 fold more abundant than the 3’ end of the gene whereas the expression of the transcript containing the 5’ part of the *mtMutS* mRNA was 1.8±0.15 fold higher than that of the extreme 3’ end of the transcript (Figure 5). This observation suggests the existence of different variants of the same gene under normal conditions in the mature mt-mRNA pool of *Sinularia* cf. *cruciata* (For details on the primers used for this assay see Additional file 4, Table 2 and Additional file 5 for results).

**Figure 5:**
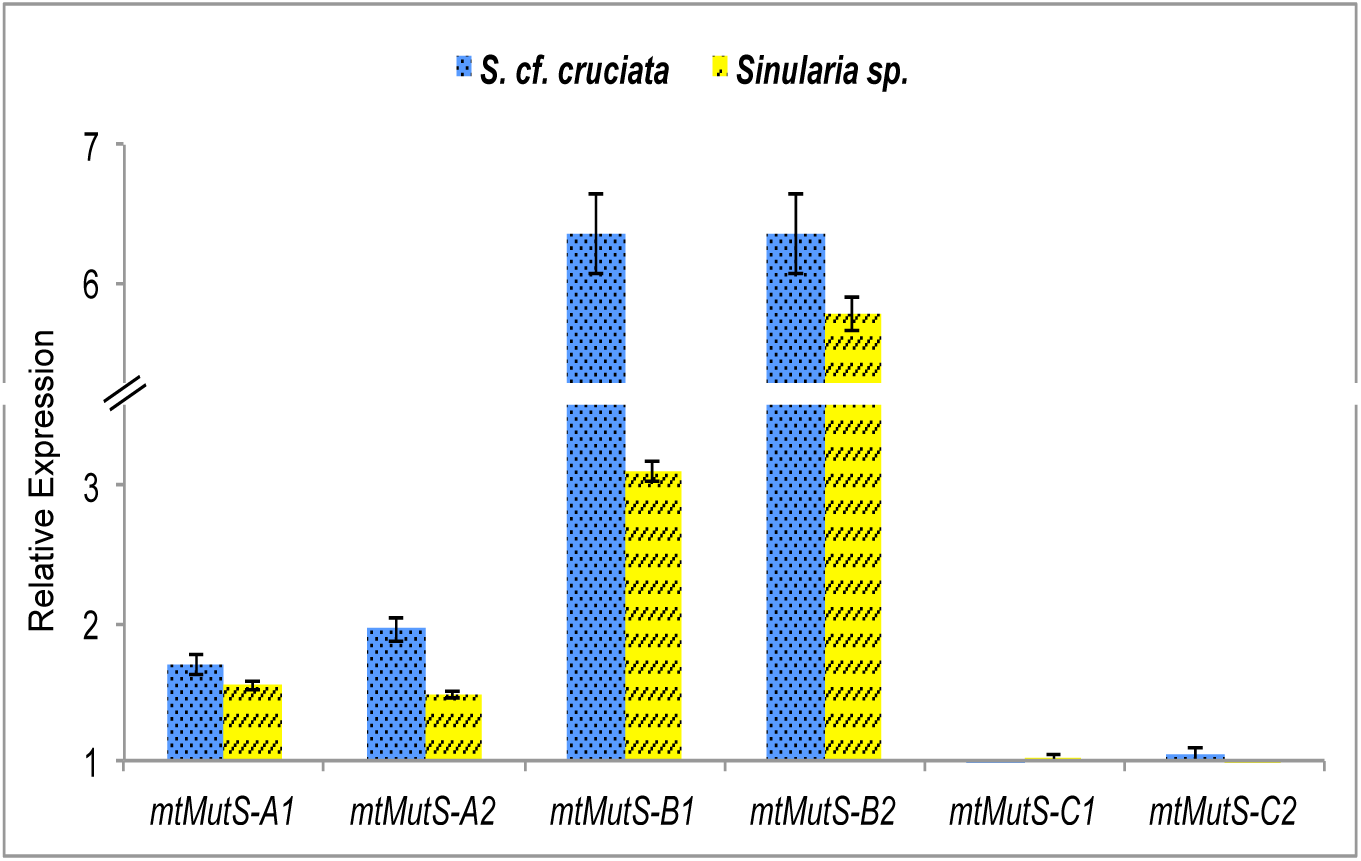
Relative quantification of alternate mRNA transcript abundance. Quantification of alternatively polyadenylated *mtMutS* transcripts. The *ACTB* gene served as reference. The comparison was performed against the 3’ region cleaved-off after RNase H digestion (*mtMutS*-C1, C2) (see Additional file 4 Table 2 for primer details). Data shows relative expression ±SD of technical triplicates for two *Sinularia* species.

### Antisense *ATP6* mRNA

Using 5’ RACE we detected antisense mRNA transcripts complementary to the *ATP6*. Five different starting points were determined for these transcripts (i.e. positions 16837, 16866, 16870, 16891, and 16920), which were longer than 200bp (reverse primer used binds the antisense messenger at position 17048), polyadenylated (i.e. reverse transcribed using anchored oligo-dT), and lacked open reading frames (ORFs). Therefore, this RNA species can be categorized as long noncoding RNA (lncRNA) [50]. Antisense strand-specific RT-PCR of an internal region of these *lncATP6* transcripts further corroborated their presence (see Additional file 6).

## 4 Discussion

Using different experimental approaches we describe, the mitogenome expression patterns of an early branching animal. The precise ends of most mature mt-mRNAs were assessed for the first time in octocorals. Most mature protein-coding mRNAs detected were bicistronic or tricistronic units, with the exception of *COI*, *ND1* and *COIII*. The occurrence of polycistronic mature mRNAs potentially stems from the paucity of tRNA punctuation marks in the octocoral mitogenome. The majority of mature transcription units were found to possess 5’ and 3’ UTRs, contrary to what is known for bilaterians in general. Moreover, the occurrence of alternative polyadenylation (APA) sites of the *mtMutS* mRNA transcript and long non-coding antisense *ATP6* RNA (*lncATP6*) provide a glimpse into a unique and potentially complex mitochondrial transcription mechanism in octocorals and likely in other early branching metazoans with non-canonical mitogenomes.

The evolution of specific mechanisms for the expression of the mitogenome was necessitated by the reduced gene repertoire and compact nature of this crucial cell organelle with its own genome evolved from an Îś-proteobacterium-like ancestor [51, 52]. In cnidarians, which contrary to a typical animal mitogenome containing 37 (13 PCGs, 2 rRNA and 22 tRNA) genes in total, harbor only 16±2 (13 or 14 PCGs, two rRNAs and one or two tRNA) genes, reduction in gene content, but not genome size, is remarkable to some extent. Our understanding of the mitogenome transcription/expression patterns and regulation is currently limited to a handful of bilaterian members of the animal kingdom [19–21]. However, such attempts are lacking for non-bilaterians. Information on the mitochondrial transcriptomes of sea anemone [53], hydrozoan [28] and jellyfish [29] have shown the potential complexity of mitochondrial transcription and highlighted the need for a better understanding of the evolutionary processes leading to different strategies of mitochondrial transcription and regulation.

The processing of the mt-tRNAs interspersed in the mitogenomes of most animals provide a mechanism to liberate protein-coding mRNAs from polycistronic precursors, leading to their maturation and availability for translation [24]. Most studies in animals so far point to the generality of the tRNA punctuation model of mitochondrial mRNA processing with only the occurrence of two bicistronic transcription units (i.e. *ATP8-ATP6* and *ND4L-ND4*) apparently due to the overlap of their ORFs [20, 21, 23]. Cnidarians, however, possess only one or two tRNA genes in their mitogenomes. Our observation of four polycistronic units comprising 11 (out of 14) different PCGs is staggering, and provides evidence for a potentially unique mechanism of mitochondrial mRNA processing, expression and regulation in octocorals. These findings may also apply to other animals exhibiting a paucity of mt-tRNA genes in their mitogenome, for example, chaetognaths [54], some demosponges [55] and other cnidarians [56].

The use of RNA-Seq data provides a unique opportunity to explore mitogenome expression in non-model organisms such as non-bilaterian metazoans. Nevertheless, we observed that very few reads mapped to the published/sequenced mitochondrial genomes (< 0.018%). This has been observed before using different NGS methods in sea anemones (0.053% reads) [53], and hydrozoans (< 0.003% reads) [28], where very few reads could be identified as mitochondrial. We assembled almost the entire mitogenome of *G. ventalina* with the exception of <3% of its nucleotides mostly belonging to IGRs flanking mature transcription units and to the *mtMutS* gene. In the case of *Corallium* and *Sinularia*, and despite the difference in the number of reads available for mapping in these two species, mapping did not result in a sufficient number of reads to produce reliable contigs. Yet, the presence of read-pairs spanning multiple genes allowed us to postulate the presence of polycistronic mt-mRNAs in these species. Emblem et.al [53] suggested that low mitogenome copy number per haploid nuclear genome results in low-level expression of mitochondrial mRNAs and is responsible for the depleted number of mitochondrial reads observed in three different sea anemones. This may explain our results as well. The poor coverage observed for the *mtMutS* gene remains puzzling. However, its absence may suggest that its function under normal physiological conditions is not required and hence its expression is kept at very low levels under such conditions.

Previous studies on octocorals have proposed the *COII-COI* IGR as potential CR/oriH in the octocorals with mitogenome arrangement “A” [30–32]. The observed absence of RNA-Seq reads in *G. ventalina* supports this proposition. The presence of very similar stem-loop structure in the *ND6-COI* IGR in *C. rubrum*, along with absence of RNA-Seq reads in this region of this species, suggests that this IGR is the potential CR in octocorals with “C” type mitogenome arrangement and corroborates earlier predictions in this regard [32, 49] (Additional file 3).

Untranslated regions (UTRs) flanking the mature mRNA transcripts play a crucial role in the post-transcriptional regulation of gene expression [57]. In mitochondria, mature mRNA transcripts are generally devoid of UTRs, or these are only few (≤3) nucleotides long [23]. The presence of 5’UTRs for transcriptional units such as *COI*, *ND3-ND4L-mtMutS*, *ND2-ND5-ND4*, and *COII-ATP8-ATP6* mRNA, and of 3’UTRs for *CytB-ND6*, *ND4*, and *COII-ATP8-ATP6* suggests a putative role of these elements in the regulation of these genes and represent, to our knowledge, the first report of the presence of long UTRs in the mature mt-mRNAs of non-bilaterians.

Different studies have detected at least five mitogenome arrangements in octocorals so far, all of which appear to preserve four conserved gene blocks. The inversion or translocation of one of these blocks at a time is proposed to have led to five different mitogenome arrangements [31, 33]. It has been suggested that the occurrence of the genes in conserved clusters is selectively advantageous, for instance, as the genes can be co-transcribed and processed in a similar way [31, 58]. However, the evidence on selection favoring a particular mitochondria gene order is sparse in cnidarians, as they exhibit high diversity of mitogenome arrangements with no sharing of gene boundaries, particularly in the subclass Hexacorallia [31]. The transcriptional units we detected encompass genes from two distinct adjacent gene blocks (e.g. the polycistronic transcription units *CytB-ND6* and *ND3-ND4L-mtMutS*), contradicting the hypothesis of co-transcription as a selective force in keeping these genes together in conserved blocks in the mitochondria of octocorals. Maintenance of synteny within the four conserved gene blocks detected so far appears to result from the lack of recombination-hotspots that promote genome rearrangements. Our results indicate that different mitogenome rearrangements detected in octocorals have different mature mt-mRNA transcript structures and transcripts and transcriptional patterns (see *Corallium vs. Sinularia-Gorgonia*). This transcriptional diversity highlights the potential complexity of mitochondrial transcription among non-bilaterians.

In this regard, assuming the tRNA punctuation model holds true, in three out of five mitogenome arrangements observed in octocorals, i.e. “A”, “C” and “D”, tRNAMet lies at the end of either H- or L- strand and its processing would liberate both coding and non-coding parts of the polycistronic precursor RNA. In *S.* cf. *cruciata*, the simultaneous or sequential processing of rRNAs could provide a mechanism for the liberation of *COI* and *ND2-ND5-ND4*. However, the excision of other transcription units from the polycistronic precursor remains to be explained. Secondary structures such as stem-loops are likely involved in maturation of pre-mRNA in octocorals, as is the case in hydrozoans and other animals [19, 28]. In *S.* cf. *cruciata*, IGRs where excision is required in order to liberate detected transcriptional units (i.e. *ND1-CytB*, *ND6-ND3*, *COIII-ATP6*) form one or more stem-loop structures (Additional file 7A). We pose that the enzymes involved in mRNA maturation recognize the conserved 11bp motifs (Additional file 7B) present in the IGRs flanking transcription units and cleave them from the precursor to be available for maturation.

The *mtMutS* gene present in octocorals is thought to underpin several peculiar processes not present in other animal mitochondria. The presence of a large gene not uninvolved in energy production, within a streamlined organelle genome dedicated to this task is mysterious as well as interesting. The second largest gene in *Sinularia* cf. *cruciata* mitogenome, *ND5* (1818bp long) is known to be the most tightly regulated protein-coding gene in other animals [59]. Thus we hypothesize that the transcription of *mtMutS* is tightly regulated as well. In favor of this hypothesis, we observed distinct *mtMutS* variants resulting from the use of different internal polyadenylation sites within the *mtMutS* gene. In contrast to its function in plant mitochondria and bacteria, which polyadenylate RNA to promote their degradation [60], polyadenylation is used in mammals to provide stability to the mature mRNA and create the stop codon, if it is not complete [26]. Polyadenylated truncated transcripts destined to degradation have also been detected in mammals [27], but, their abundance is low and they are generally difficult to detect using standard methods. All the *mtMutS* variants reported here were readily detectable indicating their potential functional role.

Interestingly, these transcripts were differentially expressed with a transcript variant encompassing Domain III and V of *mtMutS* (position 6542-8950) being more abundant under normal conditions than either the 5’ or 3’ end regions of the gene. These domains are either structurally important (e.g. Domain III) or have important biochemical functionality (e.g. ATPase; Domain V) [35, 61]. Alternative polyadenylation plays a crucial role in regulating gene expression [62]. Hence, alternative polyadenylation of the *mtMutS* gene may have a regulatory function allowing a tight control of the expression of this gene in octocoral mitochondria. The precise start and end points of each *mtMutS* mRNA variant deserved to be determined to better understand how the start-stop codons are chosen during translation. Additionally, protein studies need to be conducted in order to corroborate the localization and functionality of these transcripts and their products.

Long noncoding RNAs (lncRNAs) have been recently described in the mitochondria of mammals, primarily for *ND5*, *ND6* and *CytB*, and have been shown to interact with their mRNA complements stabilizing them and/or blocking the access of mitochondrial ribosomes, thereby inhibiting translation [63]. The presence of an lncRNA transcript for *ATP6* (*lncATP6*) in Sinularia mitochondria is striking, and indicates that the regulation of mitochondrial expression using lncRNAs evolved early in Metazoa and is ancient.

More than 99% of the mitochondrial proteome is encoded by the nuclear genome [64]. The loss of mt-tRNAs in cnidarians is suggested to have occurred in association with the loss of nuclear-encoded mt-aminoacyl-tRNA synthetases [65] and indicates a greater nuclear dependency in cnidarians relative to other animals. The retention of a single mt-tRNA for formyl-methionine is interesting and likely reflects its very specific mitochondrial function or the necessity of an excision starting point that triggers the mt-mRNA maturation cascade. Our findings, together with the general paucity of tRNAs and the varied mitogenome rearrangements observed in octocorals indicate a highly complex and perhaps a unique system for mRNA processing in the mitochondria of these organisms.

## 5 Conclusion

Recent studies on the human mitochondrial transcriptome revealed an unexpected complexity in expression, processing, and regulation of mt-mRNAs [19, 66]. Our results shed first light on the potentially more complex nature of these processes in the mitochondria of early branching animals by virtue of their “special” and diverse mitogenomes. Overall, due to the lack of tRNA punctuation marks, mitochondrial mRNA processing in octocorals appears to be drastically different. The presence of polycistronic mature mRNAs for the majority of genes provides evidence for the complexity of the transcription process in these animals. The occurrence of alternately polyadenylated transcripts for the *mtMutS* gene and their differential expression, the existence of 5’ and 3’ UTRs, and the presence of lncATP6 transcripts are additional features highlighting the diverse set of post-transcriptional modifications and regulatory mechanisms used among octocorals. More research will contribute to better understand the mitochondrial biology of early branching animals from a functional perspective. This will certainly increase our knowledge on the innovations that shaped the evolution of these organisms.

## Acknowledgements

The research was funded by DAAD (German Academic Exchange Service) PhD scholarship awarded to G.G.S. We are thankful to the editor and two anonymous reviewers of Axios Review for insightful feedback on the manuscript. We would like to acknowledge Gabi Büttner and Simone Schätzle for their assistance in the laboratory, Dr. Dirk Erpenbeck and Dr. Oliver Voigt for their constructive discussions, and Dr. Peter Naumann’s for his assistance in the aquarium. SV is indebted to M. Vargas Villalobos, S. Vargas Villalobos, S. Vargas Villalobos and N. Villalobos Trigueros for their constant support.

## Additional Files

The following Additional Files were deposited at Open Data LMU:

Additional file 1: Mitochondrial genome of *Sinularia* cf. cruciata.

Additional file 2: Mitochondrial gene expression determined by RNA-Seq for *Sinularia* cf. *cruciata*, *Gorgonia ventalina*, and *Corallium rubrum*.

Additional file 3: Stem-loop structures and conserved motifs of putative control regions from octocorals mitogenomes studied.

Additional file 4: Additional tables 1 and 2).

Additional file 5: RHAPA analysis.

Additional file 6: Antisense strand-specific RT-PCR.

Additional file 7: Stem-loop structures and conserved motif of IGRs between detected transcriptional units from *S.* cf. *cruciata*.

## Authors’ contributions

GGS, SV, and GW conceived and designed the experiments; GGS performed the experiments and analyzed the data; AP helped in bridging the mitogenome sequence gaps and analyzing the genome; GGS drafted the manuscript; and SV and GW helped with its revision. All authors read and approved the final manuscript.

